# Characterization of neural infection by Oropouche orthobunyavirus

**DOI:** 10.1101/2024.10.11.617875

**Authors:** Kaleigh A. Connors, Maris R. Pedlow, Zachary D. Frey, Jackson J. McGaughey, Gaya K. Amarasinghe, W. Paul Duprex, Leonardo D’Aiuto, Zachary P. Wills, Amy L. Hartman

## Abstract

Oropouche fever is a re-emerging global viral threat caused by infection with Oropouche orthobunyavirus (OROV). While disease is generally self-limiting, historical and recent reports of neurologic involvement highlight the importance of understanding the neuropathogenesis of OROV. In this study, we characterize viral replication kinetics in neurons and microglia derived from immortalized, primary, and induced pluripotent stem cell-derived cells, which are all permissive to infection. We demonstrate that ex vivo rat brain slice cultures can be infected by OROV and produce antiviral cytokines and chemokines, including IL-6, TNF-α and IFN-β, which introduces an additional model to study viral kinetics in the central nervous system. These findings provide additional insight into OROV neuropathogenesis and in vitro modeling strategies for a newly re-emerging arbovirus.

## Introduction

Oropouche orthobunyavirus (OROV) is a newly re-emerging arbovirus endemic to South America, (1). Belonging to the *Peribunyaviridae* family, OROV is transmitted to humans primarily through the bite of the *Culicoides paraensis* midge and potentially mosquitos such as *Culex quinquefasciatus, Aedes aegypti, Ochlerotatus serratus* (2,3). Since its identification in Trinidad in 1955, OROV has infected tens of thousands of individuals primarily in South America (4,5), where it remained largely under the radar compared to the more well-known Dengue, Chikungunya, and Zika viruses. The disease Oropouche fever is generally a febrile illness with severe headache, chills, arthralgia, myalgia, and maculopular rash. Infrequently, cases of hemorrhagic fever and neurological involvement have been noted (2,6,7), and until recently, human deaths have not been associated with OROV infection. At the end of 2023, larger outbreaks of OROV began in both endemic and new areas of South America (8,9). Cuba reported its first ever locally acquired case in June of 2024, and there have been several travel-associated cases in the United States and Europe. As of September 2024, 9,852 confirmed cases of Oropouche fever have been reported in Central and South America, the United States, and Canada, with Brazil reporting the largest number of cases. Most concerningly, this recent outbreak has seen higher levels of apparent severe disease including neurological issues, development of Guillain-Barre Syndrome (GBS) (10), and the first ever reported deaths due to hemorrhagic manifestations of OROV disease in Brazil (11). Alarmingly, cases of mother-to-child-transmission, resulting in microcephaly and even fetal death, have been reported for the first time (12). It is currently unknown whether the apparent severity of this most recent outbreak is due to circulation of a more pathogenic strain of OROV or other factors. Some early evidence indicates that the currently circulating strain is a result of a reassortment event and that it may perhaps also have faster replication kinetics in vitro (13).

Little is known about OROV neuroinvasive disease, which is estimated to occur in about 4% of cases (6,10,14,15). Neurologic symptoms - dizziness, photophobia, confusion, nystagmus - raise concerns about the virus’s ability to breach the blood-brain barrier and cause direct neuronal damage. The mechanisms by which OROV may invade the central nervous system (CNS) are not well understood, hindering the development of effective therapeutic interventions.

To better understand potential neuropathogenesis of OROV, we characterized OROV infection in several model systems, including immortalized human and rodent cell lines, primary cells, human induced pluripotent stem cell-derived (hiPSC-derived) neurons, and ex vivo rat brain slice cultures (BSCs). Neural progenitor cells (NPCs), neurons, and microglia from different species were all highly permissive to OROV infection, resulting in virus amplification and cell death. Moreover, OROV infection in ex vivo BSC induces inflammatory cytokines including IFN-β, IL-1β, and MCP-1. These findings provide tools for studying OROV infection in the CNS and begin to address major gaps in understanding the neurovirulence of this re-emerging virus of public health importance.

## Materials & Methods

### Biosafety

Work with OROV was completed in a Biosafety Level 2 (BSL-2) laboratory following all university biosafety guidelines.

### Animal work

All work with animals adhered to The Guide for the Care and Use of Laboratory Animals published by the NIH throughout the duration of the study. The University of Pittsburgh is fully accredited by the Association for Assessment and Accreditation of Laboratory Animal Care (AAALAC). The University of Pittsburgh Institutional Animal Care and Use Committee (IACUC) oversaw this work and approved it under protocol numbers 21100165 and 22051190.

### Viruses

The BeAn19991 strain of OROV was rescued through reverse genetics and was generously provided by Paul Duprex and Natasha Tilston-Lunel (Pitt Center for Vaccine Research) (16). Virus was propagated in Vero E6 cells with standard culture conditions using Dulbecco’s modified Eagle’s medium (DMEM) (ATCC, 30-2002) supplemented with 1% penicillin/streptomycin (Pen/Strep), 1% l-glutamine (l-Glut), and either 2% (D2) fetal bovine serum (FBS). A standard viral plaque assay (VPA) was used to determine the titer of the stocks. The agar overlay for the VPA was comprised of 2X minimal essential medium (MEM, ThermoFisher, 11935046), 2% FBS, 1% Pen/Strep, 1% HEPES buffer, and 0.8% SeaKem agarose (Lonza, BMA50010); the assay incubated at 37°C for 4 days, followed by visualization of plaques with 0.1% crystal violet.

### Cells

All BV2 cells (provided by Gaya Amarasinghe) and Vero cells (American Type Culture Collection [ATCC], CRL-1586) were cultured in DMEM supplemented with 1% Pen/Strep, 1% l-Glut, and either 2% (D2), 10% (D10), or 12% (D12) FBS. SH-SY5Y (ATCC, CRL-2266) were cultured in D12/F12 media (ATCC, 30-2006) supplemented with 1% Pen/Strep and 1% l-Glut. N2a (provided by Gaya Amarasinghe) and HMC-3 (ATCC, CRL-3304) cells were maintained in Eagle’s minimum essential medium (EMEM) (ATCC, 30-2003) with 10% FBS and supplemented with 1% Pen/Strep and 1% l-Glut.

Prior to infection, N2a, SH-SY5Y, BV2 and HMC-3 cells were plated in 24-well plates, on poly-L-lysine (R&D Systems, 3438-100-01) coated coverslips, except for BV2 cells where poly-L-lysine was not used. Cells were plated at the following densities: N2a 100k-200k cells/well; Sh-Sy5y 500k-1 million (mil) cells/well; BV2 75k-100k cells/well; HMC-3 100k-200k cells/well.

### Isolation and culture of primary rat cortical neurons

On the day prior to neuron isolation, acid-washed coverslips were coated with PDL/Laminin (Sigma, P7405-5MG; Invitrogen, 23017-015). Dissociation media (DM) comprised of Hanks’ Balanced Salt Solution (Invitrogen, 14175-103) supplemented with 10 mM anhydrous MgCl2 (Sigma, M8266), 10 mM HEPES (Sigma, H3375), and 1 mM kynurenic acid was prepared. DM was brought to a pH of 7.2 and sterile filtered prior to use. On the day of isolation, a trypsin solution containing a few crystals of cysteine (Sigma, C7352), 10 milliliter (mL) of DM, 4 microliter (μl) 1N NaOH, and 200 units of Papain (Worthington, LS003126); and a trypsin inhibitor solution containing 25 mL DM, 0.25g trypsin inhibitor (Fisher, NC9931428), and 10 μl 1N NaOH were prepared, and filter sterilized. Embryonic day 18 Long Evans (Crl:LE; Charles River Laboratories, Wilmington, MA) rats were dissected, and brains were removed. The cortices were separated from the hippocampus and placed into DM. 5 mL of trypsin solution was added and cortices were placed in a 37°C water bath for 4 min, swirling occasionally to mix. The trypsin solution was removed, and cortices were immediately washed with trypsin inhibitor once, and then twice more while swirling in the water bath. Following the washes, the trypsin inhibitor was removed and replaced with 5 mL of Neurobasal/B27 media, then triturated to dissociate the neurons. Final volume was brought to 10 mL of Neurobasal Plus/B27 Plus media, and cells were counted and plated at a density of 50k cells/well for 96-well plates, 100-150k cells/well for 24-well plates, or 2mil cells/well for 6-well plates. One hour after isolation, the media was removed and replaced with fresh Neurobasal/B27 media. Primary neuron cultures were maintained in Neurobasal/B27 media, which consists of standard Neurobasal Plus Medium (Gibco, A3582901) supplemented with 1% Pen/Strep, 1% L-Glut, and 2% B27 Plus Supplement (Gibco, A3582801).

### Generation of human iPSC-derived neural progenitor cells and neurons

Human NPCs were generated as previously described (17). Briefly, hiPSCs were cultured in mTeSR1-plus medium supplemented with dual SMAD inhibitors SB 431542 and LDN 193189 to promote neural induction. After 8–10 days, neural rosettes were manually isolated, transferred into Matrigel coated plates and cultured in StemDiff Neural Progenitor Medium (STEMCELL Technologies, 05833) for the expansion of NPCs. All cells were cultured in standard conditions (37°C, 5% CO2, and 100% humidity).

NPCs were seeded into Matrigel-coated 12-or 6-well plates to the density of 2.5 × 10^5^ or 5 × 10^5^ cells/well, respectively, and cultured in neurobasal medium [Neurobasal medium supplemented with 0.5X B27 (Vitamin A+), 1% Pen/Strep, 1% Glutamax, BDNF (10 ng/ml), CHIR99021 (3 μM), Dorsomorphin (1 μM), Forskolin, and ROCK inhibitor (10 μM)]. Two days later, CHIR99021, Dorsomorphin, Forskolin (10 μM), and ROCK inhibitor were withdrawn and differentiating NPCs were cultured for 4 weeks. Half medium was changed every other day.

### Ex vivo brain slice cultures

Rat brain slice cultures were generated as previously described (18). Time-mated Sprague-Dawley rats (Hsd:Sprague Dawley SD) used in our study were obtained from Envigo. Litters of > 6 pups were obtained at postnatal day six for BSC isolation. The sex of animals for BSC generation was dependent on the litter. The number of slices obtained per experiment was dependent on litter size. Briefly, Prior to slicing, 6-well plates were seeded with 10% heat-inactivated fetal bovine serum (FBS; Corning, 35-011-CV) in Neurobasal-A (Gibco, 10888022) (plating medium) and incubated at 37C, 5% CO2. Whole brains were isolated from 6-day-old rat pups following decapitation. The cerebellum was cut with a straight razor to yield a flat coronal surface on the posterior aspect of the brain, which was then affixed vertically to the vibratome stage with glue. Ice-cold dissection media (1X PBS, 5 mg/ml D-glucose, 1% Penicillin-Streptomycin) was used in the chamber, which was kept cold using ice cubes. Progressive, 400 μm thick coronal slices were obtained at speed 8, frequency 9, after the bi-lobular frontal lobe was removed. Between 3-5 slices were obtained per brain. Slices were transferred to a petri dish with cold-dissection media. Slices were then transferred to lie flat on a hydrophilic PTFE membrane (EMD Millipore, PICM03050) placed in each well of a 6-well plate using a metal spatula. After 24 hours, inserts and slices were transferred to new 6-well plates containing Neurobasal-A with 5% FBS. After 72 hours, inserts and slices were transferred to new 6-well plates containing Neurobasal-A with 1x B27-A (Neurobasal-A/B27 media) (Gibco 12587010). Media was replaced every other day until 10 days in vitro (DIV) when slices were infected with 1 × 10^5^ pfu OROV per slice.

### Viral infection

Cells were plated in 12 or 24-well plates as indicated above prior to infection. Virus was diluted to indicated MOIs in 100-200 μl D2 media. Complete culture media was removed from wells and replaced with viral inoculum or mock (D2 only) infected. Cells were incubated for 1 hour at 37°C, 5% CO2 with rocking every 10-15 min. For immortalized cell lines, inoculum was removed and cells were washed once in 1X PBS, then cultured in D2 media. For primary neurons and iPSC-derived human NPCs and neurons, viral inoculum was removed and replaced with complete culture media. Supernatants were collected at indicated timepoints for vRNA in Trizol reagent (Invitrogen, 15596026) or frozen at -80°C for VPA.

Rat brain slices were inoculated by pipetting 200 μL inoculum onto the top of the slice and 800 μL around the PTFE insert. Plates were incubated at 37oC, 5% CO2 for 1 hour adsorption. In the meantime, new 6-well plates were seeded with 1 mL Neurobasal-A/B27 media. Using forceps, PTFE membranes were transferred into the fresh 6-well plates for culture. Viral replication was obtained by quantifying infectious titer from the supernatant collected at 24, 48 and 72 hpi using VPA. Release of lactate dehydrogenase (LDH) from damaged cells was assessed per manufacturer’s instructions using LDH Glo Cytotoxicity Assay (Promega, J2381) as previously described (18).

### RT-qPCR

RNA isolation was performed with an Invitrogen PureLink RNA kit (Invitrogen, 12-183-025). Samples were inactivated 1:10 in Trizol reagent. 200 μl of chloroform was then added to each sample and allowed to sit at room temperature for three minutes. The samples were then centrifuged for 15 min at 12,000 x g at 4°C. After separation of the aqueous phase, the aqueous phase was removed, and an equivalent volume of 70% ethanol was added. The remainder of the isolation was performed with the PureLink RNA protocol. qRT-PCR was then performed using the Invitrogen SuperScript III Platinum One-Step Quantitative Kit (Invitrogen, 11732020). OROV primers targeting the S segment used were: OROV19991-Forward 5’-TACCCAGATGCGATCACCAA-3’ and OROV19991-Reverse 5’-TTGCGTCACCATCATTCCAA-3’. The Taqman probe used was OROV19991-Probe 5’-6-FAM/TGCCTTTGGCTGAGGTAAAGGGCTG/BHQ_1-3’. qRT-PCR was performed on the Quantstudio 6 (Applied Biosystems).

For cytokine/chemokine measurements, rat BSCs were homogenized in 200 μL Neurobasal-A/B27 media through agitation with a pipet. Total RNA was extracted from 100 μL of BSC homogenate using the RNAeasy mini kit (Invitrogen, 12183025) with DNAse treatment. cDNA was synthesized using M-MLV reverse transcriptase (Invitrogen, #28025013) with random hexamer primers. Semi-quantitative real-time PCR was used as previously described (18,19) for the detection of cytokines and chemokines (IFNα, IFNβ, IL-1β, IL-6, TN-α, MCP-1/CCL2).

### Antibodies

The following primary antibodies were used for immunocytochemistry staining at 1:500 dilution: rabbit anti-OROV N (Custom Genescript), mouse anti-OROV serum (in house), anti-mouse Nestin (EMD-Millipore, MAB5326), anti-chicken Nestin (Novus Biologicals, NB100-1604), anti-chicken Beta III tubulin (EMD Millipore, AB9354). Secondary antibodies were used at 1:500 dilution: goat anti-mouse AF488 (Invitrogen, A11001), goat anti-mouse AF647 (Invitrogen, A21235), goat anti-rabbit AF594 (Invitrogen, A11012), goat anti-rabbit AF488 (Invitrogen, A11008), goat anti-chicken AF594 (Invitrogen, A32759) or goat anti-chicken AF647 (Invitrogen, A32933TR).

### Immunofluorescence

Upon harvest, virus-infected cells were fixed in 4% paraformaldehyde for 15 min, followed by 3 washes in 1X PBS. Coverglass was permeabilized with 0.1% Triton X-100 detergent in 1X PBS for 15 min at room temperature (RT). Cells were blocked using 5% normal goat serum (NGS) (ThermoFisher, 50062Z) for 1-3 hours (h) at RT, followed by incubation with primary antibodies for 1-2 h at RT. After 3 washes in 0.5X NGS, the secondary antibodies were added for 1 h at RT. The cells were counterstained with Hoescht 33258 (Invitrogen, #H1398, 1:1000) and mounted using Gelvatol. Fluorescent slides were imaged using a Leica DMI8 inverted fluorescent microscope provided by the Center for Vaccine Research at 20X magnification or using a Nikon A1 confocal fluorescent microscope at the Center for Biologic Imaging. Images were processed and quantified using Fiji (20).

### Statistical analysis

Statistical analyses were performed using GraphPad Prism software (La Jolla, CA). For Figure 3B and 3C, a one-way analysis of variance (ANOVA) was performed to compare results between OROV infected BSCs to mock infected controls across timepoints. Multiple comparison was performed using Brown-Forsythe test. Significance indicated by: *, P<0.05; **, P<0.01; ***, P<0.001; ****, P<0.0001; ns, no significance.

## Results

### Oropouche orthobunyavirus replicates in neurons and microglia

To characterize OROV infection in resident brain cells, we assessed replication kinetics in CNS-related immortalized cell lines derived from mice and humans. Undifferentiated murine N2a cells were infected with OROV at an MOI of 0.1 or 0.01. Infection resulted in a 3-log increase in viral RNA titers (as measured by plaque forming unit equivalents/ml; pfu eq./ml) by 72 hours post infection (hpi) (**Figure 1A**). Undifferentiated human neuroblastoma SH-SY5Y cells were infected with OROV at an MOI of 0.01 or 0.001 where we also observed a 3-log increase in viral titers by 72 hpi (**Figure 1B**). Immunofluorescent microscopy of viral antigen at 48 hpi show substantial staining of both N2a and Sh-SY5Y cells alongside antibodies against Pax6 (neural progenitor cells), Nestin (neural progenitor cells), and Beta III tubulin (neuronal differentiation) (**Figure 1C**).

**Figure 1.**
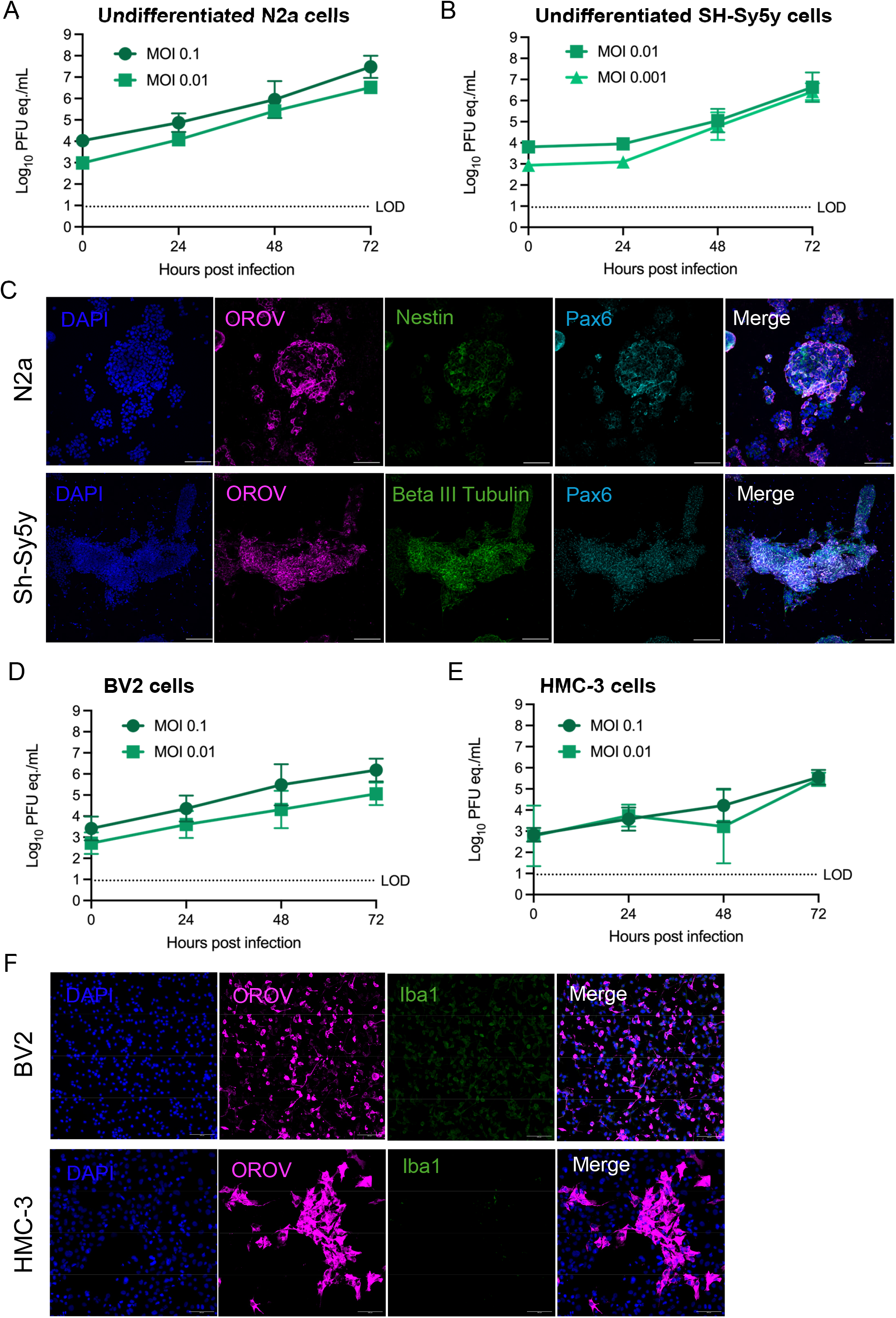
Oropouche orthobunyavirus infects and replicates in neural cell lines in a dose-dependent manner. Neuronal (A-C) or microglial (D-F) infection with OROV. (A) Undifferentiated murine N2a cells, (B) undifferentiated human SH-SY5Y, (D) murine BV2, and (E) human HMC-3 cells were inoculated with OROV at MOI 0.1, 0.01, or 0.001 and viral RNA was quantified at 24, 48 and 72 hpi by RT-qPCR. The data for each cell line comprises two separate experiments. (C) N2a and SH-SY5Y were fixed at 48 hpi and immunostained for anti-OROV (magenta), anti-Nestin (green, top panel), anti-beta III tubulin (green, bottom panel), anti-Pax6 (cyan) and counterstained with DAPI (blue). (F) BV2 and HMC-3 cells were fixed at 48 hpi and immunostained for anti-OROV (magenta), anti-IBA-1 (green), and counterstained with DAPI (blue). Slides were imaged at 20x magnification. Scale bar = 100 μm.

We then assessed OROV replication kinetics in microglial cell lines. Both BV2 (mouse) cells and HMC-3 (human) cells were infected with OROV at an MOI of 0.1 or 0.01. Viral RNA titers increased by 3-logs in BV2 cells and 2-logs in HMC-3 cells by 72 hpi (**Figure 1D-E)**. Immunofluorescent microscopy of BV2 and HMC-3 cells demonstrates widespread viral antigen staining by 48 hpi (**Figure 1F**). These findings demonstrate that immortalized neurons and microglia are highly permissive to OROV in a dose dependent manner.

### Primary neural progenitors and neurons are permissive to OROV infection

Next, we assessed OROV replication kinetics in primary rat cortical neuron cultures and human induced pluripotent stem cell-derived (hiPSC-derived) neuroprogenitor cells (NPCs) and neurons. Primary neuronal cells more accurately represent the morphological and physiological state of cells in vivo. Primary rat neurons infected with MOI 0.1, 0.01, or 0.001 OROV resulted in infectious titers up to 1 × 10^5^ pfu/ml by 48 hpi (**Figure 2A**). Microscopy of neurons infected at MOI 0.1 revealed diffuse viral antigen staining at 24 hpi (**Figure 2B**).

**Figure 2.**
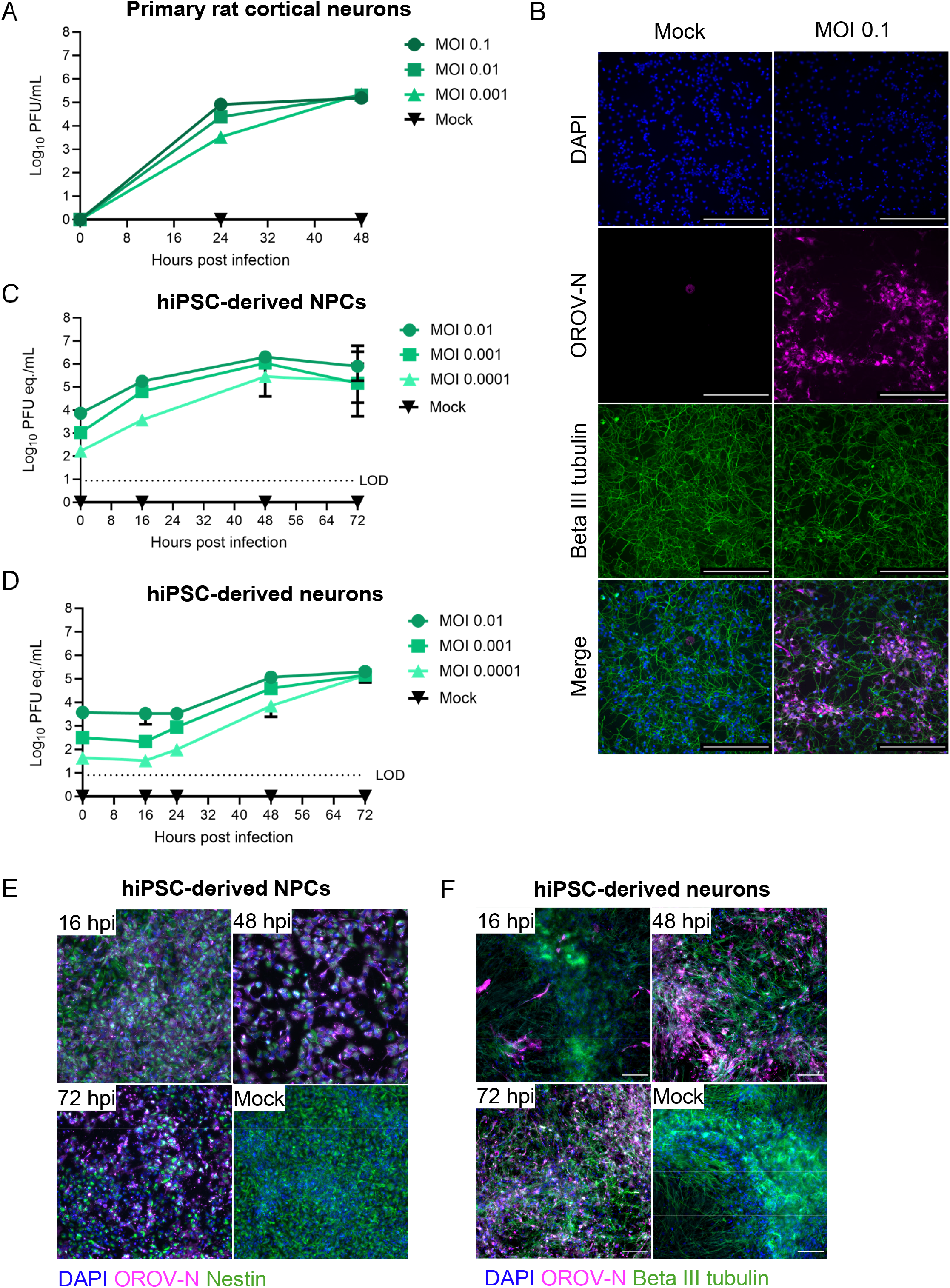
Oropouche orthobunyavirus replicates to high titers in neural progenitor cells and neuronal cultures. (A) Primary rat cortical neurons were infected at 4 days in vitro with OROV at MOI 0.1, 0.01, or 0.001. Viral plaque assay was used to quantitate infectious virus at 24 and 48 hpi. (B) Primary rat cortical neurons were fixed at 24 hpi and immunostained with anti-OROV-N (magenta), anti-beta III tubulin (green), and counterstained with DAPI (blue). Slides were imaged at 20x magnification. Scale bar = 250 μm. Human iPSC-derived NPCs (C) and neurons (D) were inoculated with OROV at MOI 0.01, 0.001, or 0.0001. Viral RNA was quantitated at 16, 24, 48 and 72 hpi by RT-qPCR. (E) hiPSC-derived NPCs and (F) hiPSC-derived neurons were fixed at 16, 48 and 72 hpi alongside mock-infected cells and immunostained with anti-OROV-N (magenta), anti-Nestin (E, green) or anti-beta III tubulin (F, green) and counterstained with DAPI (blue). Slides were imaged at 20x magnification. Scale bar = 100 μm.

We then used human iPSC-derived NPCs and neuron cultures (**Figure 2C-E**). Human iPSC-derived NPC cultures retain the ability to differentiate into various cell types, allowing for comparison of viral permissivity between immature and mature neuronal states (21). A similar dose-dependent replication was observed in hiPSC-derived NPC and neuron cultures over 72 hpi, with a 2-log increase in viral titers in NPCs at 48 hpi (**Figure 2C**), and 1.5-log increase in neuronal culture by 72 hpi (**Figure 2D**). Interestingly, there seems to be differential kinetics of virus replication in the two culture types. In NPCs, OROV replicates with an immediate increase in vRNA and antigen staining by 16 hpi. In the neuronal cultures, there was an apparent lag time where increases in viral titer were not detected until 48 hpi. This can be observed using immunofluorescent miscroscopy, were the viral antigen staining of NPCs at 16 hpi is diffuse (**Figure 2E**), while in the neuron culture there are only a few positive cells observed until later time points (**Figure 2F**). These results demonstrate that primary neurons and NPCs from different species are highly permissive to OROV infection, where the virus replicates to high titers.

### OROV infection in brain slice culture induces pro-inflammatory cytokines

Ex vivo brain slice culture (BSC) preserves structural integrity of neurons and resident cells of the CNS, including astrocytes and microglia, which play pivotal roles in viral infection (22). We previously established and characterized rat BSC as a model for Rift Valley fever virus (RVFV), a related neurovirulent bunyavirus (18). Using this same model, we characterized OROV growth kinetics and immune response to infection. Individual coronal slices from postnatal day 6 rats were inoculated with 1 × 10^5^ pfu OROV, and supernatant was collected up to 72 hpi. Infectious viral titer, measured by plaque assay, indicated that ex vivo rat BSC supported viral replication over time (**Figure 3A**), with peak titer reaching 1 × 10^3.5^ pfu/mL at 48 hpi. As a proxy for tissue damage and cell death, we found increased release of lactate dehydrogenase (LDH) into supernatants collected from OROV-infected slices compared to mock infected controls (**Figure 3B**), suggesting virus-induced tissue damage as a result of OROV infection.

**Figure 3.**
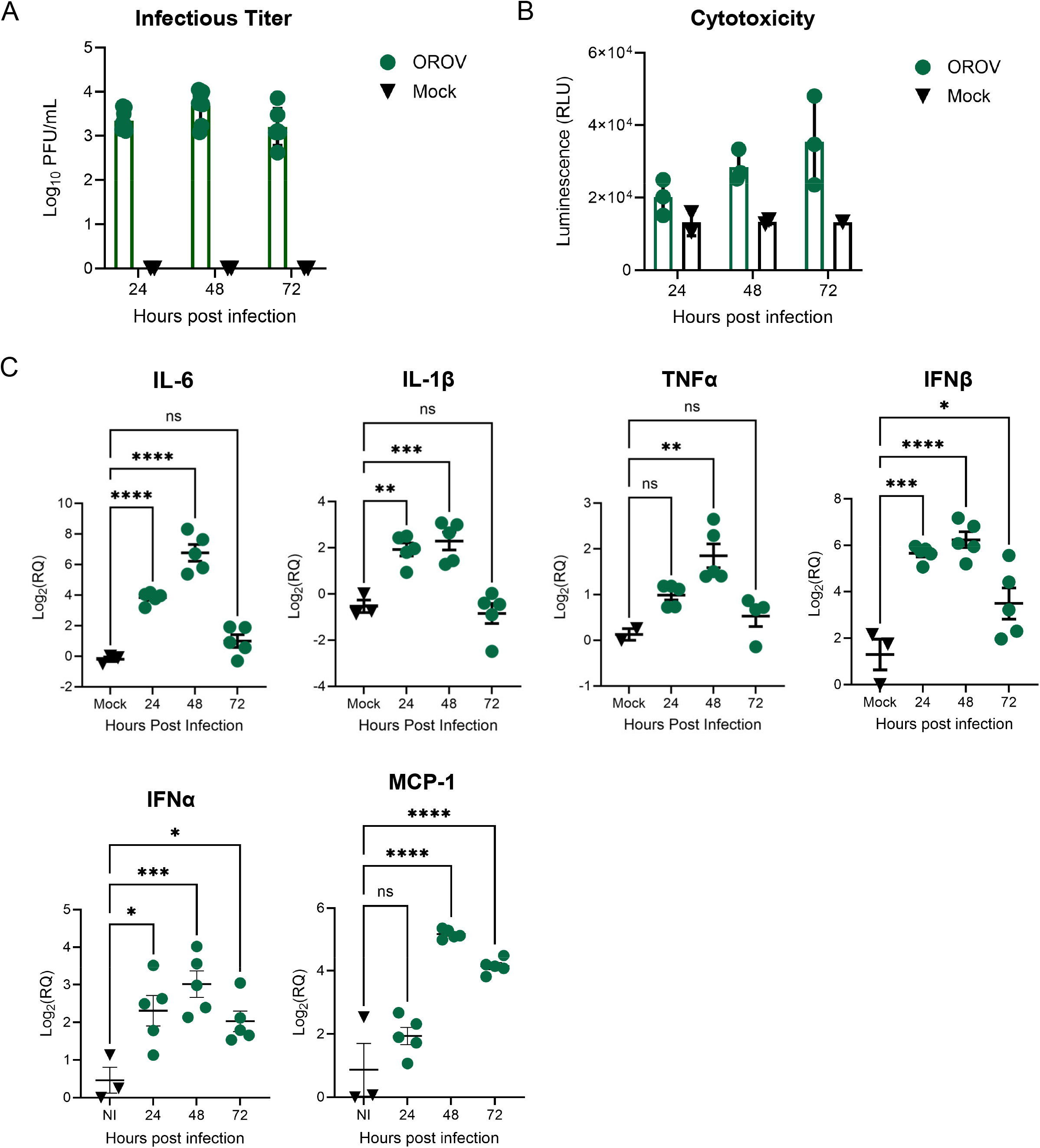
OROV infects and replicates in ex vivo rat brain slice cultures and induces an antiviral response. (A) Coronal rat brain slices were inoculated with 1 × 10^5^ pfu OROV or mock infected. Infectious titer from supernatants were quantitated using viral plaque assay at 24, 48 and 72 hpi. Data is a combination of OROV-infected brain slices (n = 8) and mock-infected brain slices (n = 4) across two independent experiments. (B) Cytotoxicity was quantified using LDH-Glo assay at 24, 48 and 72 hpi from supernatant collected from OROV-infected (n = 3) or mock infected (n = 2) slices. (C) Rat BSC were infected with 1 × 10^5^ pfu OROV (n = 5) or mock infected (n = 3) and RNA was isolated at 24, 48 or 72 hpi from whole slice. Mock data represents mock infected slices from each timepoint, and gene expression data was pooled. Gene expression was quantified by RT-qPCR using delta-delta Ct and normalized to the variation of the amount of beta-actin within each slice. Fold changes are relative to a 0-hour mock infected control slice. Error bars represent SEM. Statistical analysis performed using a one-way ANOVA. *P < 0.05; **P < 0.01; ***P < 0.001; ****P < 0.0001; ns = not significant.

To get a basic understanding of the inflammatory response to OROV infection in BSC, we measured changes in cytokine and chemokine gene expression over time in homogenized BSC samples. By 24 hpi, there were significant increases in the expression of IL-6, IL-1β, IFN-β, and IFN-α in BSCs infected with OROV compared to the mock-infected control slices (**Figure 3C**). At 48 hpi, a significant increase in gene expression is observed for all cytokines and chemokines tested: IL-6, IL-1β TNF-α, IFN-β, IFN-α and MCP-1. These findings support the notion that OROV induces a pro-inflammatory response from resident neural cells in response to infection.

## Discussion

While OROV is primarily known for causing Oropouche fever, a self-limiting febrile illness, growing evidence suggests that the virus may have neuroinvasive properties, leading to CNS involvement (6,14). In fact, OROV infection has been linked to miscarriage, stillbirths, and microcephaly in infants, which may previously have been reported but not further explored (12,23). For the first time, OROV infection has resulted in the development of GBS in adults (10). This aspect of OROV pathogenesis has been relatively underexplored despite its potential implications for public health.

A limited number of animal models for OROV pathogenesis exist. In 1961, Anderson et al. demonstrated that immunocompetent Swiss-Webster mice inoculated intraperitoneally (i.p.) show no clinical signs, while 2-day-old suckling mice succumb to disease in 2-3 days (24). In contrast, intracranial (i.c.) inoculation in immunocompetent mice is lethal, regardless of age (24). Rodrigues et al. demonstrated a 30% lethality in a golden hamster (*Mesocrisetus auratus*) model of OROV infection, with an abundant amount of viral antigen detected the brain and liver of animals by histopathology (25). Santos et al. showed that subcutaneous dorso-lumbar inoculation of suckling BALB/c mice resulted in 85% lethality, with the central nervous system being the main target of infection, potentially from the brainstem to cerebral cortex (26,27). While immunocompetent adult mice are generally resistant to OROV infection, immunocompromised mice have been used to determine that susceptibility to lethal OROV infection is dependent on mitochondrial antiviral signal protein (MAVS) (28). Further analysis demonstrates that in mice lacking a complete type I interferon signaling system, OROV infection concentrates in the liver, spleen and blood (28,29). Interestingly, deletion of interferon regulatory factor 5 (IRF-5) results in encephalitic disease and delayed lethality, highlighting the role of IFNs in controlling peripheral infection and mediating neuroinvasion in mice (29).

Prior reports from mouse models indicate both neurons and microglia are susceptible to OROV infection (25,26,30). Indeed, we demonstrate that immortalized neurons and microglia, primary neurons, and human iPSC-derived neural cells are permissive to OROV infection. Neural cells obtained from both rodent (mouse and rat) and human origin support viral replication in a dose-dependent manner. Interestingly, we found higher levels of vRNA in mouse neurons when compared directly to the mouse microglia cell line BV2. This may be due to the fact that microglia are more immunoreactive, and more readily induce an antiviral immune response, potentially limiting OROV replication in a monolayer culture system (31). The role that microglia play during OROV infection, including potential harmful effects of microglial activation, is an area in need of further study.

Human NPCs were highly susceptible to OROV infection with a faster replication cycle, even at low MOIs, compared with hiPSC-derived neuronal cultures and primary rat neurons. NPCs are found in the developing and adult nervous system and can proliferate and differentiate into different types of neural cells, including neurons, astrocytes, and oligodendrocytes (32). NPCs retain significant plasticity, allowing them to respond to environmental signals and generate specific cell types needed for CNS development, repair, and maintenance. Further, NPCs play a critical role in neurogenesis during embryonic development and contribute to neural plasticity and repair in the adult brain. NPC infection by many viruses, including Zika virus (ZIKV), Japanese encephalitis virus (JEV), and Lymphocytic Choriomeningitis virus (LCMV), results in premature differentiation, reduced proliferation, dysregulation of cell cycle, and cell death (33–37). The effects of OROV infection in neurogenesis requires further evaluation.

Ex vivo BSCs offer an additional tool to understand neural susceptibility to viral infection and local immune response to viral infection through maintenance of cytoarchitecture and neural connections (22,38). A prior study found that human brain slices were susceptible to OROV infection, with a preference for microglia, followed by neurons, although there was no astrocyte infection (30). This is supported with an in vivo study that indicates astrocytes are activated during OROV infection of neonatal mice without noting infection of astrocyte cells (26). Here, we show that rat BSCs are susceptible to OROV infection resulting in the production of infectious virus and accumulation of tissue damage over time. Additionally, rat BSCs infected with OROV produce an antiviral immune response, characterized by increased in antiviral and inflammatory cytokines TNF-α, IFN-β, and IFN-α, which is in line with previous reports (39,40). We observed significant increases in antiviral cytokine and chemokine expression, albeit to lower levels than was induced by RVFV in ex vivo rat BSCs (18). This highlights the differences in neurovirulence, and potential differences in virulence factors between OROV and RVFV. This study did not assess viral tropism in the rat BSCs; however, these analyses are planned for future studies.

The strain of OROV used in these analyses, BeAn1991, was isolated from a sloth in Brazil in 1991 and falls within Lineage I (41). This is also the strain used to develop the OROV reverse genetics system (16). The 2024 outbreak in Brazil is the result of a recombination event leading to a new OROV lineage, BR-2015-2023, which may have enhanced replication and immune evasion allowing it to emerge (42). Future studies are necessary to understand if the currently circulating strain has enhanced neuroinvasion and/or neurovirulence capacity in comparison with BeAn1991.

Experimental models are needed to support study of emerging and re-emerging viruses, including for OROV. This report describes in vitro and ex vivo models for studying OROV pathogenesis in the CNS and the antiviral immune response to infection. Our results indicate permissivity of neural progenitor cells to OROV infection, which is critical to explore given the recent cases of vertical transmission and the described microcephaly. Additional investigations are necessary to understand the molecular and immune mechanisms which support OROV infection in the CNS.

## Acknowledgements

This work was funded in part through National Institutes of Health (NIH) grants R56 AI171920, R01 NS101100, R01 AI178378, R01 AI169850. The authors declare no competing interests exist.

